# Amortized template-matching of molecular conformations from cryo-electron microscopy images using simulation-based inference

**DOI:** 10.1101/2024.07.23.604154

**Authors:** Lars Dingeldein, David Silva-Sánchez, Luke Evans, Edoardo D’Imprima, Nikolaus Grigorieff, Roberto Covino, Pilar Cossio

## Abstract

Biomolecules undergo conformational changes to perform their function. Cryo-electron microscopy (cryo-EM) can capture snapshots of biomolecules in various conformations. However, these images are noisy and display the molecule in unknown orientations, making it difficult to separate conformational differences from differences due to noise or projection directions. Here, we introduce cryo-EM simulation-based inference (cryoSBI) to infer the conformations of biomolecules and the uncertainties associated with the inference from individual cryo-EM images. CryoSBI builds on simulation-based inference, a combination of physics-based simulations and probabilistic deep learning, allowing us to use Bayesian inference even when likelihoods are too expensive to calculate. We begin with an ensemble of conformations, which can be templates from molecular simulations or modelling, and use them as structural hypotheses. We train a neural network approximating the Bayesian posterior using simulated images from these templates, and then use it to accurately infer the conformations of biomolecules from experimental images. Training is only done once, and after that, it takes just a few milliseconds to make inference on an image, making cryoSBI suitable for arbitrarily large datasets. CryoSBI eliminates the need to estimate particle pose and imaging parameters, significantly enhancing the computational speed in comparison to explicit likelihood methods. We illustrate and benchmark cryoSBI on synthetic data and showcase its promise on experimental single-particle cryo-EM data.

## 1 Introduction

Biomolecules constantly reorganize their structure to perform their biological functions. Channels and transporters cycle between open and closed conformations to regulate traffic through cellular membranes. Molecular motors alternate between different states to advance in discrete steps. A pathogen’s invasion protein—like the spike of SARS-CoV-2— opens up to anchor to the surface of our cells to infect them. Understanding the mechanisms underlying biomolecular function requires the knowledge of their alternative structures and an understanding of how they interconvert.

An accurate determination of the various important molecular conformations is challenging. Many experimental techniques report average observables. Single-molecule methods can report on dynamics but are limited in their structural resolution. On the computational side, molecular dynamics (MD) simulations provide trajectories at high temporal and spatial resolution. However, the inability to sample relevant timescales and force-field accuracy issues limit simulation approaches. Integrative methods in structural biology combine experimental and computational techniques with the promise to describe a more complete picture of biomolecular phenomena [1].

Cryo-electron microscopy (cryo-EM) enables the study of biomolecular structures at atomic resolution [2, 3]. In cryo-EM, a transmission electron microscope records two-dimensional projected images of a thin sample containing many molecules (micrographs). Specialized software then cuts out the two-dimensional images containing single molecules (particles) [4]. The sample is prepared by flash-freezing an aqueous solution of randomly oriented biomolecules. Since freezing is very fast [5, 6, 7], the biomolecules are trapped in all of their different conformations. To avoid radiation damage, the frozen sample is imaged with a limited number of electrons, resulting in picked particles that are very noisy projections of the molecule’s density in unknown conformations and orientations.

Cryo-EM 3D classification methods can identify different particle conformations in a dataset, provided the dataset contains a sufficient number of particles in each conformation [8]. Reconstructing a molecular conformation at high resolution requires averaging over many different particles that depict the same conformation. Most 3D classification methods assume that a dataset contains particles that can be classified into a relatively small set of conformational states. The classes are then iteratively refined by explicitly comparing multiple poses (*i*.*e*., rotations and translations) of the current best class averages to the experimental particle using a cross-correlation or least squares metric within a likelihood calculation [9, 10, 11] to minimize inter-class variability. The 3D maps can be successfully combined with MD approaches to extract a conformational ensemble consistent with each cryo-EM map [12, 13, 14].

However, rare conformations, as well as conformations that are structurally similar but functionally different, might be missed in such an analysis. This could lead to an incomplete analysis of the full landscape of conformations needed to understand the molecule’s biological function [15]. To obtain high-resolution reconstructions, each 3D class has to have a sufficient number of particles to cover the entire projection (*i*.*e*., reciprocal) space. Particles belonging to classes with insufficient sampling and other low-resolution classes are often discarded. Moreover, the refinement of classes often does not converge and repetitions of the same analysis with different random seeds leads to particles assigned to different classes [16]. These problems make it difficult to identify rare conformations or transition states, which occur far less frequently and might therefore be missed.

Machine learning (ML) has enabled an important step towards extracting conformational heterogeneity from cryo-EM. Milestones in the field were set by manifold embedding [17] and deep generative models [18, 19, 20, 21, 22] to represent conformational landscapes. The key idea is to use ML to learn the mapping between the particles in a cryo-EM dataset and the corresponding conformational volumes [23]. However, the statistical inference can become computationally intractable, especially when both the conformation and pose are inferred simultaneously. This is why most cryo-EM ML methods, as well as non-ML variability methods [24, 25, 26, 27], rely on costly explicit-likelihood methods to calculate the particle poses. cryoAI [28] and its implementation in cryoDRGN [29] use direct gradient-based optimization to amortize the particle poses while still requiring the direct estimation of a pose for each particle. Furthermore, these methods do not obtain likelihood estimates or statistical errors in assigning each particle to a given conformation.

Template-matching based approaches assign a particle to a molecular structure with high fidelity. The BioEM approach can be used to discriminate molecular conformations from individual particles by integrating over poses and imaging parameters within a Bayesian framework [30, 31]. Recently, high-resolution 2D template matching has been applied to identify biomolecular conformations *in situ* using a cross-correlation-based approach [32, 33]. When the poses are sampled on a fine angular grid, and the matching is repeated for templates representing different molecular conformations, this method provides a highly accurate metric for conformational identification. However, this brute-force likelihood optimization becomes quickly intractable.

Here, we leverage recent advances in simulation-based inference in combination with molecular simulations to develop cryoSBI, a computational framework that matches a particle image to molecular three-dimensional conformations without having to search the template poses or calculate any likelihood explicitly. Additionally, our framework provides amortized inference and accurate statistical confidence. Learning high-resolution structures and conformational variability simultaneously from a cryo-EM is a highly challenging problem. We, therefore, decompose the problem in two steps. First, we assume that one (or a few) high-resolution structures can be reconstructed using conventional methods. Second, we assume that molecular simulations such as advanced MD schemes [34] or ML tools such as AlphaFold [35] can provide a set of conformations that serve as structural hypotheses starting from one (or few) experimental structures. We then simulate cryo-EM experiments to produce template particles that answer the question: “What would an experimental particle look like if this was the molecular conformation?”. The synthetic particles arise from random poses and imaging parameters with adequate noise levels. Neural network density estimation allows us to learn from these particles the Bayesian posterior, which quantifies the most probable structure corresponding to a particle. We illustrate the cryoSBI algorithm, benchmark it on synthetic data, and showcase its use on experimental particles.

## 2 Results

### 2.1 Simulation-based inference of single-particle cryo-EM

We formulate the task of inferring molecular conformations from single cryo-EM particle image as a Bayesian inference problem. In essence, we wish to quantify the probability that a given image *I* depicts a molecular conformation *X*. Let us consider a set of molecular conformations parametrized by a vector *θ*. In the following, we will assume without loss of generality that *θ* is one dimensional, *i*.*e*., a function *f* (*X*) = *θ* that maps molecular configurations to a real number. Given a cryo-EM image *I*, we aim to infer the conformation *θ* of the molecule observed in the image. Therefore, we want to compute the Bayesian posterior *p*(*θ*| *I*), quantifying how compatible *θ* (that is associated with conformation *X*) is with the observed image *I*. The posterior can be computed by Bayes’ theorem

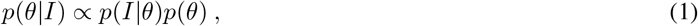

where *p*(*I*|*θ*) is the likelihood of generating an image *I* given a molecular conformation *θ* and the prior *p*(*θ*) encodes all available knowledge before making the inference on how the conformations are distributed. Modeling the image formation process described by the likelihood requires taking into account the details of the experiment and the functioning of the electron microscope. For instance, that the molecule is randomly rotated, that the image will be noisy, and that the microscope will introduce an aberration. The full likelihood of the image processing is, therefore, *p*(*I*| *θ, ϕ*), which contains the additional parameters *ϕ* required to model the details of the experiments and microscope. Usually, we are not interested in making inference of *ϕ*, which are therefore nuisance parameters. In other words, the likelihood is actually a marginalization

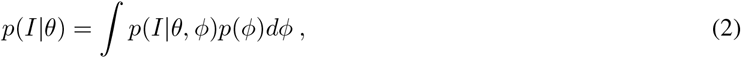

with the prior of the imaging parameters *p*(*ϕ*). Maximizing the marginal likelihood *p*(*I*|*θ*) is computationally very expensive [31, 32] because each evaluation requires integrating over all imaging parameters. The high computational cost of approaches that require explicit likelihood optimization restricts their applicability to small sets of images.

Simulation-based inference (SBI) is an alternative approach to do Bayesian inference with intractable or computationally expensive likelihoods [36, 37, 38, 39]. The main idea is to replace a likelihood evaluation with a forward model to simulate synthetic data and then learn an approximation of the posterior on them. Here, we develop an SBI framework for amortized template-matching of conformations from single-particle cryo-EM images (Figure1). While the inverse problem of inferring a conformation from a cryo-EM image *I* is challenging, the forward problem is much simpler. We can easily encode the image formation process described by the likelihood *p*(*I* |*θ, ϕ*) in a cryo-EM simulator, and repeatedly sample it by running forward simulations to produce synthetic images, *i*.*e*., *I*_*i*_ ∼*p*(*I*| *θ*_*i*_, *ϕ*_*i*_) with *θ*_*i*_ ∼*p*(*θ*), *ϕ*_*i*_ ∼*p*(*θ*). In this way, we accumulate a data set of images and parameters describing their generation, *D* = *{θ*_*i*_, *ϕ*_*i*_, *I*_*i*_*}*. We used Neural Posterior Estimation[40] to directly approximate the Bayesian posterior from the data set of images *D*. We first used an embedding network *S*(*I*) to extract features and map each image into a medium-dimensional representation. We then used another neural network *q* as a conditional density estimator to build a surrogate model of the posterior, *i*.*e*., a statistical model that approximates the posterior, *q*(*θ*| *S*(*I*)) ≈ *p*(*θ*| *I*). We then trained *S* and *q* jointly on *D* using standard supervised deep learning methods. In this way, we learned an approximation of the desired Bayesian posterior, bypassing any explicit likelihood evaluation and marginalization.

**Figure 1:**
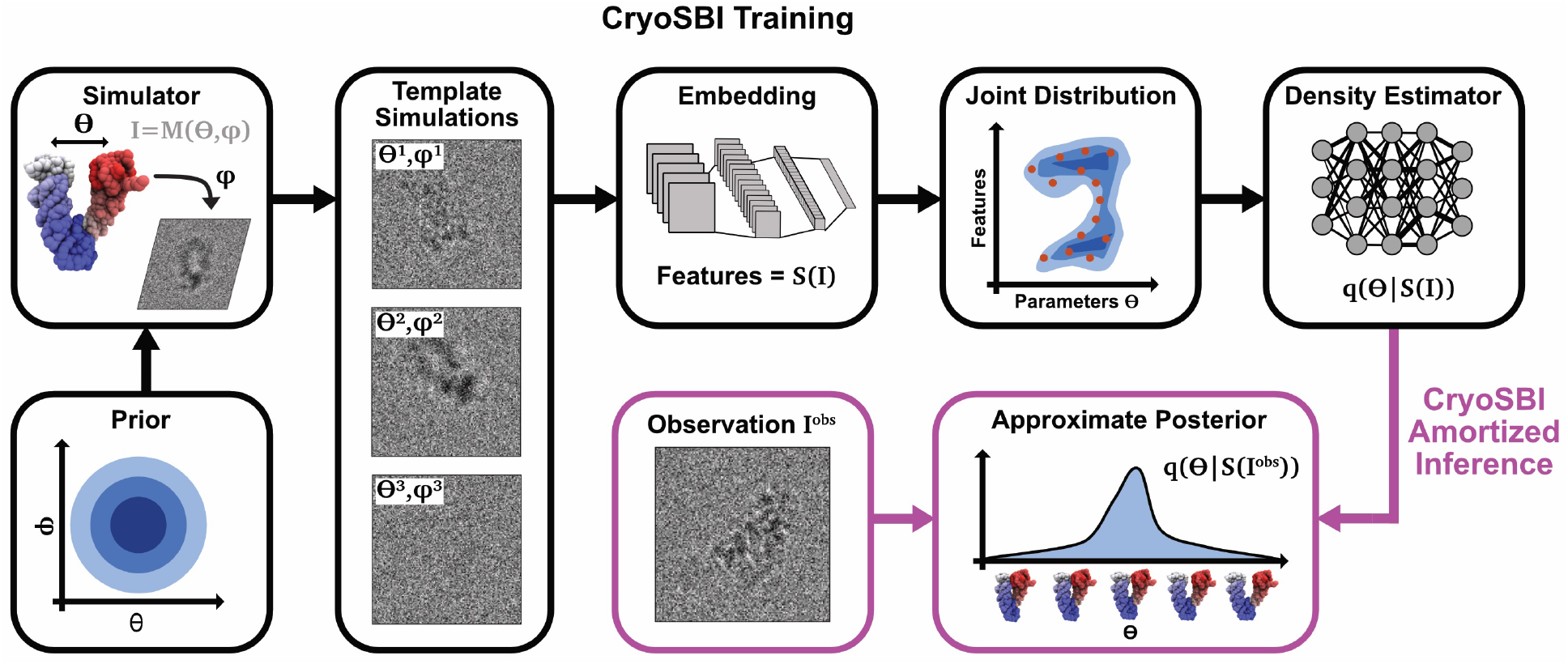
Schematic representation of cryoSBI: Simulation-based inference for cryo-EM. Simulated particle image templates generated from conformations *θ* and imaging parameters *ϕ*, such as rotation, defocus, and translation, are used to train the cryoSBI embedding network and posterior estimator (black boxes). *θ* and *ϕ* are sampled from a prior distribution, and fed to a forward model *M* to generate a particle image *I*. An embedding network that featurizes the images *S*(*I*) and a density estimator of the joint distribution of features and parameters are trained simultaneously using millions of simulated particle images. CryoSBI learns a computationally efficient approximation of the posterior *p*(*θ*| *S*(*I*)), which enables amortized template matching of experimental images (magenta boxes). For each experimental particle, cryoSBI provides the full posterior, indicating both the most probable conformation that generated the image and the statistical confidence of the inference.

After training on the simulated images, the neural density estimator *q* estimates the posterior for any new experimental image. The inference is computationally efficient for two reasons: first, it does not require any marginalization over the nuisance parameters *ϕ*; and second, the inference is amortized. The computationally expensive part due to the sampling must be paid only once upfront by repeatedly running the simulator to build the data set *D*. Once the conditional estimator *q* is trained, any new inference requires only an evaluation of the neural network underlying *q*.

In summary, cryoSBI involves the following steps:

1. Prepare a data-set of experimental cryo-EM particles obtained with conventional workflows.
2. Obtain a set of molecular conformations that serve as structural hypotheses—template structures—by using molecular simulations or ML methods.
3. Simulate many synthetic particles sampling all possible template structures and nuisance parameters.
4. Obtain a surrogate of the Bayesian posterior by training the embedding and conditional density estimator simultaneously on the set of simulated template particles.
5. Perform inference on the experimental particles with the trained surrogate posterior.

### 2.2 Validation and benchmark with synthetic data

How precisely is it possible to identify a structure in a single cryo-EM image? We answered this question by validating and benchmarking cryoSBI using synthetic data obtained from Hsp90, an established benchmark model in the field [41, 42]. Hsp90 comprises two chains that perform a large conformational change corresponding to their opening and closing. We selected twenty structures spanning the opening of the chains, measured by the root-mean-squared-deviation (RMSD) with respect to the closed structure (Figure 2A).

**Figure 2:**
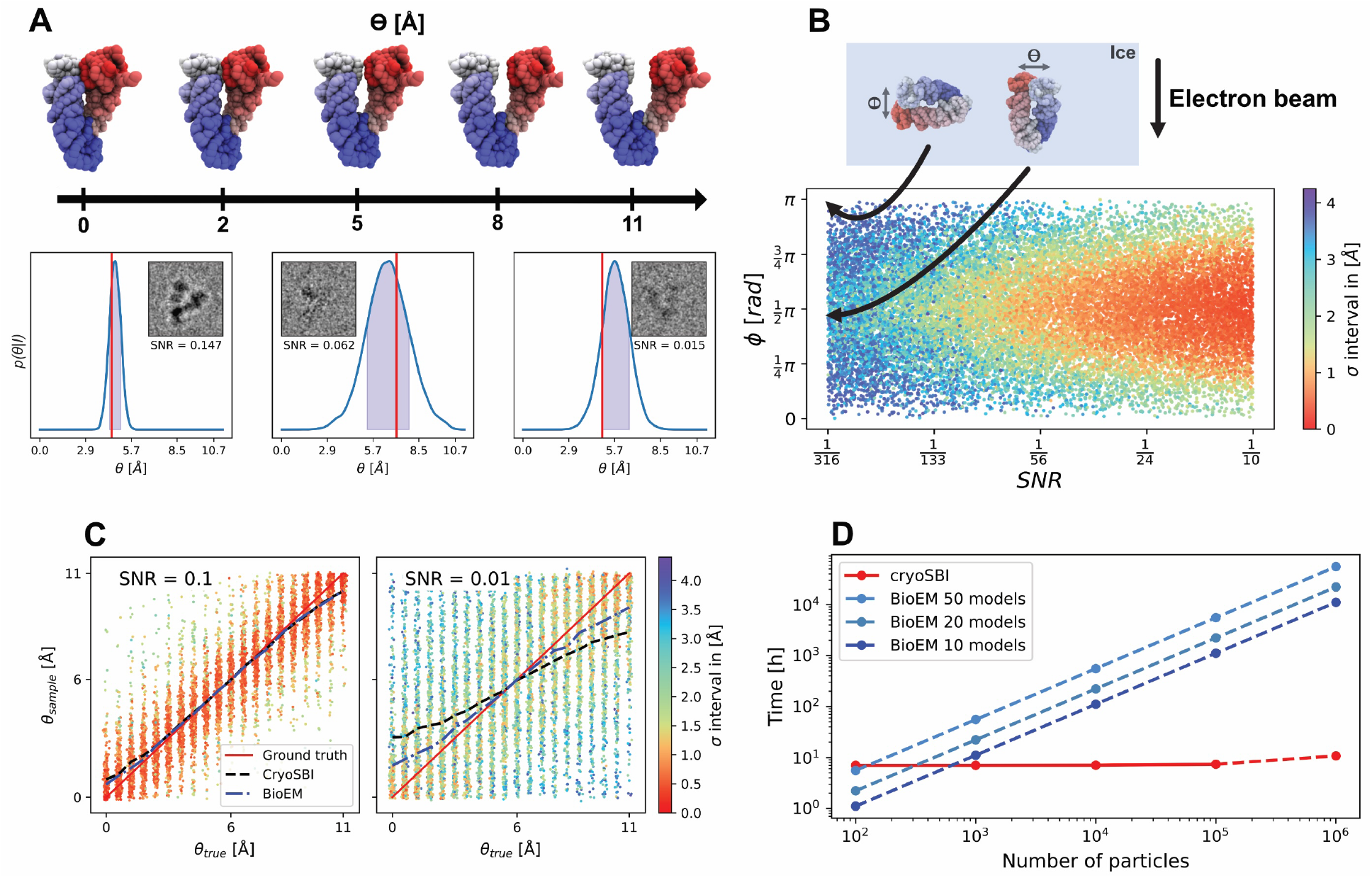
Validation and benchmark with synthetic data. A) The opening of the two arms of Hsp90 defines a conformational change, quantified by *θ*, the RMSD w.r.t the closed conformation. We selected 20 configurations equally spaced along *θ* as structural models. Examples of cryoSBI inference for three particles. Each inference on synthetic images (gray-scale images in the insets) resulted in a posterior (blue curve), quantifying which structural model was the most compatible with the image. Red lines represent the true configurations that generated the images in the insets. The shaded region is the 2*σ* interval. B) Inference precision for 10,000 images as a function of SNR and projection direction. Each point represents the *σ* interval (color bar, from dark blue - low confidence, to deep red - high confidence) of the posterior obtained from a synthetic image produced at a given SNR and projection angle. On top, we show the schematic definition of *ϕ*, the angle formed between the direction of the electron beam and the direction of movement of the two arms of Hsp90. C) Inference accuracy for 10,000 images for sets with different SNR. The two scatter plots show the correlation between the estimated opening of Hsp90 obtained as a sample of the posterior, and the true opening, using SNR=0.1 and SNR=0.01, respectively. We drew one sample from the posterior for each image, and colored them according to the posterior *σ* interval (color palette, colors as for panel B). The black dashed line describes the average of the posterior means, while the blue line corresponds to the mean of the maximum-likelihood estimates obtained via BioEM [31]. D) Scaling of the computational cost w.r.t. the number of images of cryoSBI and maximum likelihood methods (BioEM). The computational cost corresponds to wall-time using an NVIDIA RTX A6000 GPU for cryoSBI, and AMD EPYC 7742 CPU for BioEM.

CryoSBI accurately infers molecular configurations from single images. We trained cryoSBI on synthetic cryo-EM particle images generated under realistic conditions, with random orientations, a wide range of defocus values, center translation, and signal-to-noise ratio (SNR). We learned a surrogate model of the posterior, that we used to make inference on the selected synthetic particles. For each inference, we obtained an estimate of the Bayesian posterior that we could compare with the structure that we actually used to produce the specific image (red line in Figure 2A). The posterior provides information both on the accuracy of the inference—whether the bulk of the distribution contains the ground truth—and the precision—the spread of the distribution.

The SNR and projection direction are the main experimental factors determining how precisely we can infer a molecular configuration from a single image. As the SNR of an image decreases, the inference is still accurate but the posterior gradually broadens, corresponding to an increasing uncertainty (Figure 2A,B). The precision also decreases for projection directions that “hide” the conformational change of interest. In the case of Hsp90, this occurs by projecting along a direction where one arm covers the other one (Figure 2B), where the projection direction is parallel to the relevant conformational motion. For very low SNR and bad projection directions, the inference returns an approximately flat posterior. In other words, cryoSBI correctly tells us that these images cannot be reliably assigned to specific configurations.

A more systematic evaluation confirms that cryoSBI is accurate and precise for SNRs within experimental range [41]. We compared inferred configurations to ground truths for 10,000 images, assuming high and low SNR (Figure 2C). For SNR=0.1, 68% of inferred configurations were accurate within 1 Å with an average uncertainty of 0.75Å measured by the posterior *σ* interval (color bar). For SNR=0.01, the accuracy of the prediction declined slightly to 68% of the inferred structures being within 2.7 Å of the true structure. The average uncertainties rose to 2.1Å. In comparison to an explicit likelihood method (dashed lines), cryoSBI’s predictions are slightly worse for low SNR. This is expected, as SBI methods tend to lose some accuracy due to the approximations of the ML models that make amortization possible, a phenomenon also observed in 3D reconstruction [43]. We note that for flat posteriors, the mean is biased toward the center due to cryoSBI being trained on a finite domain of *θ*, therefore, we used samples of the posterior that do not exhibit this issue.

CryoSBI is fast and enables inference of very large data image sets and molecular conformational ensembles. We compared the computational cost of performing inference on the synthetic data set generated with Hsp90 with cryoSBI and methods that optimize an explicit likelihood model [30, 31]. These require to evaluate the likelihood and marginalize over all model parameters for each image, leading to a linear scaling of the computational cost with the number of images and the number of conformations (Figure 2D). The cost becomes quickly prohibitive (several days for 100,000 images, 20 structures using 36864 orientations on a CPU node). CryoSBI’s inference is instead amortized, that is, the largest computational cost occurs upfront to produce simulations and train the model (approximately a few hours), after which the inference is effectively for free. Each inference only requires a forward pass of the trained neural network that serves as a surrogate model of the posterior, which takes approximately a millisecond. The benefit of amortization is that the computational cost of doing inference does not scale significantly with the number of images (Figure 2D). Amortization opens the door to comparing hundreds-to-thousands of structures to datasets containing an arbitrary large number of images.

### 2.2 Validating with experimental data

Having validated CryoSBI on synthetic data, we sought to demonstrate that it is also accurate in making inferences from actual experimental particles. So far, the training templates and the synthetic data have been generated with the same forward model simulator and parameter distributions. This will not be the case with experimental data, for which we must generally assume model misspecification. In other words, the data we simulate to train the model and the experimental particles that we want to make inferences on will always differ. However, we cannot access a ground truth with experimental particles, and therefore first validate our method using a standard experimental benchmark system: apoferritin.

Apoferritin is 474 kDa large cytosolic globular protein complex composed of 24 subunits forming a hollow nanocage (Figure 3A). It is highly symmetric and rigid, making it a standard benchmark in the cryo-EM field. We used a published data set containing 483 particles of apoferritin. Due to the absence of conformational flexibility, the particles do not include alternative conformations. We selected the PDB structure built from the cryo-EM map reconstructed from the same dataset [44] as our ground truth (Figure 3A). We generated a hypothetical conformational ensemble by varying our ground truth structure along two normal modes (Figure 3A). The order parameter is *θ* = *γ* RMSD to ground truth, where *γ* =− 1 for normal mode 1 and *γ* = 1 for normal mode 10, which quantifies the distance of the resulting structures from the ground truth reference that sits at *θ* = 0 by construction. We selected normal modes 1 and 10 to ensure two distinct conformational changes, thereby avoiding degeneracies caused by symmetry.

**Figure 3:**
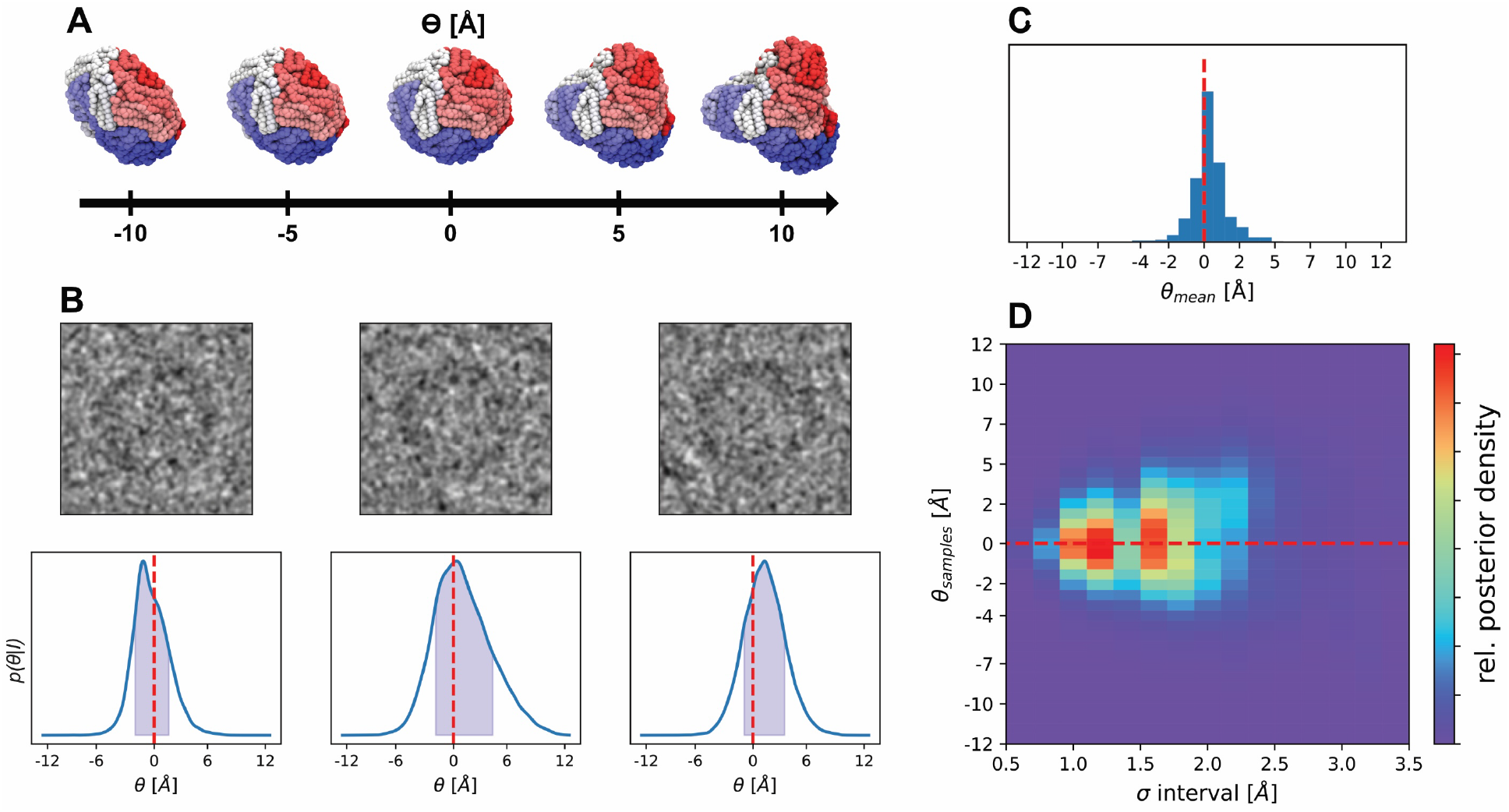
CryoSBI applied to experimental particles of apoferritin. A) Apoferritin conformational change along *θ*, generated by varying the ground truth reference cryo-EM structure (PDB ID 4v1w) along two normal mode directions. The conformational change is quantified by the RMSD in Å to the reference cryo-EM structure. We multiplied the RMSD by -1 for variations along mode 1 (left half of the axis). By construction *θ* = 0 Å for the reference structure. B) Example of cryoSBI posteriors for experimental apoferritin particle images. The particles are low pass filtered for better visibility. The red dashed line indicates the position of the reference structure along *θ*. C) Histogram of the inferred conformation from the posterior mean *θ*_mean_ for the particles in the dataset. D) Two-dimensional histogram reporting 1000 posterior samples from the posterior of each apoferritin particle.

CryoSBI correctly maps individual experimental particles of apoferritin to their corresponding reference structure. Following our pipeline, we trained a surrogate posterior on synthetic templates generated starting from the structural ensemble shown in Figure 3A. Posteriors are peaked around *θ* = 0, both for individual particles (Figure 3B) and for the entire experimental data set (Figure 3C), indicating that we could identify the 3D structure corresponding to the individual particle accurately.

The posterior width shows that the mapping is also quite precise, with uncertainty in the order of a few Angstroms. Since we are analyzing singe-molecule data (snapshots of a single protein in a specific conformation) and not averaged observables (*e*.*g*., 3D maps), we should not expect that every posterior conditioned on each image is sharply peaked precisely at the reference structure. Some particles will be more informative than others. Building a histogram by resampling each posterior conditioned on each image provides a statistical view of how informative the particles in the data set are. Interestingly, the histogram is shaped like a funnel. Particles whose posterior samples are closest to the cryo-EM structure (i.e., centered around *θ* = 0) are those that we can map with the highest confidence (Figure 3D).

### 2.4 A challenging experimental data set

Next, we challenged cryoSBI using an experimental dataset containing particles of hemagglutinin, a homotrimeric protein complex found on the surface of influenza viruses [45]. Compared to apoferritin, the hemagglutinin data set presents several additional challenges. Hemmaglutinin is more dynamic, and the particles capture a much more heterogeneous structural ensemble. In fact, only around 47% of the particles led to the reconstruction of the published high-resolution structure. Additionally, the protein adopted a preferred orientation in the experimental sample, leading to a particle distribution that does not cover uniformly the entire space of possible rotations. This can be problematic for mapping two-dimensional projections into a three-dimensional structure.

Despite these challenges, cryoSBI could correctly identify hemagglutinin configurations in single particles. We generated a hypothetical structural ensemble by perturbing the PDB structure (PDB ID 6wxb), from a 2.9Å resolution cryo-EM map [45], along two normal modes (Figure 4A-bottom), where *θ* is defined as the RMSD distance from the PDB structure. We used these structures to produce synthetic templates and train a posterior model to evaluate experimental particles. The posterior could identify the reconstructed cryo-EM structure accurately and precisely (Figure 4A). Evaluating the posterior on the entire dataset, we found a large concentration of particles (∼ 50%) that we could map to the high-resolution hemagglutinin structure with high confidence, consistent with the 47% used to generate the cryo-EM reference structure (Figure 4B). These particles are distributed in a region spanning approximately 4Å around the high-resolution structure, consistent with thermal fluctuation at room temperature, as shown by atomistic MD simulations in explicit solvent (Supplementary Figure 1). As expected, no other conformation is identified in the dataset with high confidence. For many particles, the posterior is approximately uniform and therefore uninformative.

**Figure 4:**
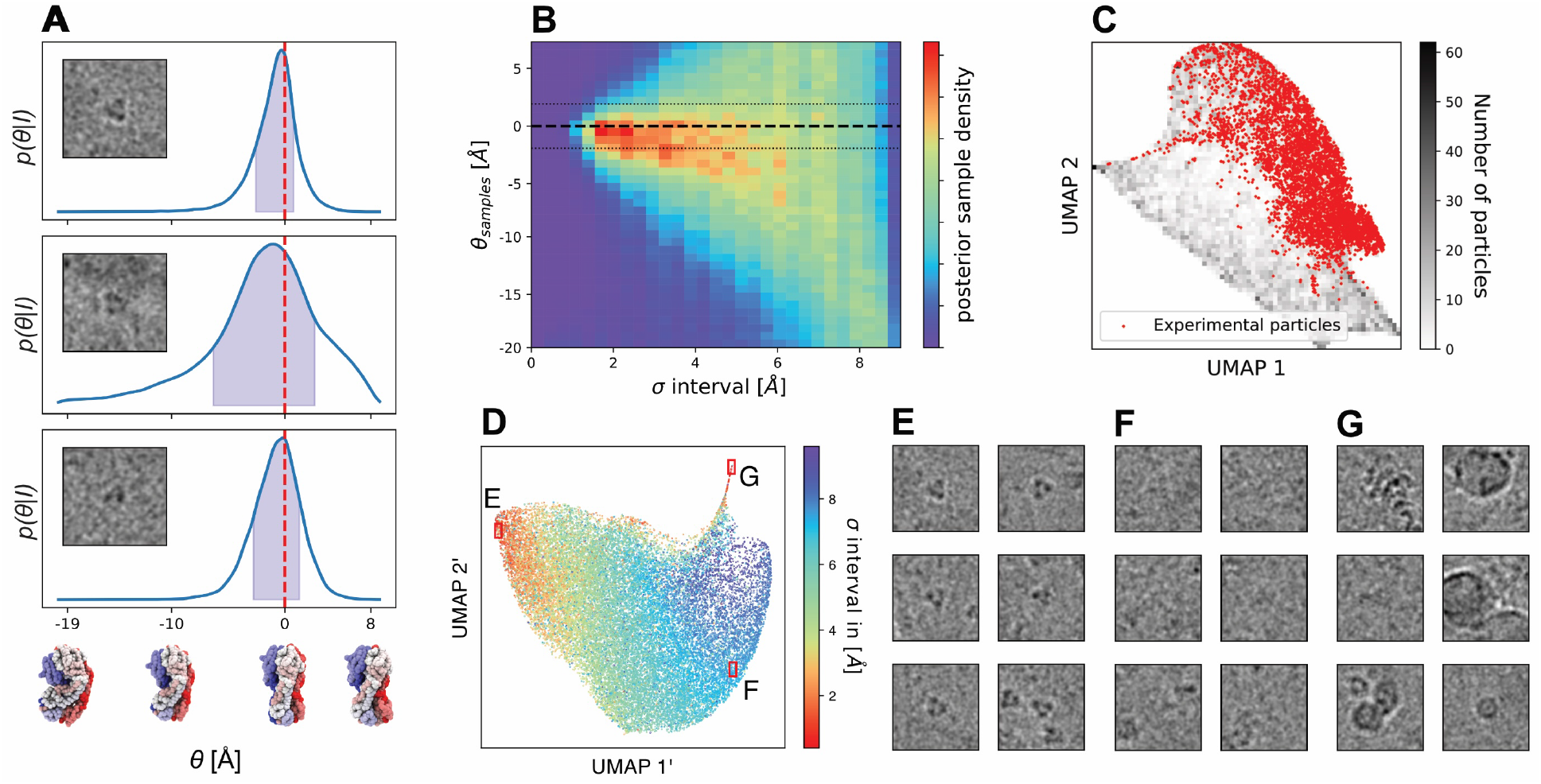
CryoSBI applied to experimental particles of hemagglutinin. A) Example of cryoSBI posteriors for experimental hemagglutinin particle images (low pass filtered for better visibility). The conformational change along *θ* was modelled normal mode analysis, *θ* = *γ* RMSD from reference cryo-EM structure (red dashed line) where *γ* =− 1 or 1 for normal mode 1 and 2, respectively. B) Two-dimensional histogram showing 1000 posterior samples for each of the 271558 posteriors. The black dashed line indicates the cryo-EM reference structure. The dotted lines show the size of the expected fluctuation around the reference structure estimated with MD simulations. C) UMAP 2D projection of the latent space of the simulated particles used for training (gray) and a random subset of 10% of experimental particles (red). D) UMAP 2D projection of the latent representation of a random subset of 10% of experimental particles colored according to their posterior *σ* interval. (E-F) Experimental particles, selected from different positions in the UMAP in D. (low pass filtered for better visibility).

### 2.5 Validating inference with latent space analysis

The embedding provides a powerful tool for analyzing the particles. The embedding is done by a neural network that encodes all particles from a 128^2^ pixels into a 256-dimensional representation. Conventional dimensionality reduction techniques can further reduce the dimensionality of the representation to generate plots that allow us to visually inspect the entire dataset. Each red point in Figure 4C corresponds to a single experimental particle, whereas synthetic particles, much more numerous, are represented as a grayscale heat map. The two coordinates, UMAP1 and UMAP2, are non-linear functions of the pixels defining the original images, which should quantify some “essential” features. Indeed, we can correlate different values of these coordinates to different modelling and imaging parameters, like the SNR, or conformations (Supplementary Figure 2).

When examining the embedding space, an initial question is whether the synthetic hemagglutinin particles used to train our posterior are consistent with the experimental particles. Figure 4C shows that synthetic particles are distributed in a region that contains the experimental ones. In other words, the simulator generates templates very similar to the experimental ones. Synthetic particles populate a larger region than the experimental ones. This is expected, and means that not all configurations, or imaging parameters, we considered as template hypotheses correspond to particles that constitute the ensemble captured by the cryo-EM experiment. A more quantitative statistical analysis based on the maximum mean discrepancy metric confirms that synthetic and experimental particles are very similar (Methods and Supplementary Figure 3). This analysis is essential to detect model misspecification, which occurs when the posterior is trained with synthetic data that do not accurately mimick the experimental data, leading to incorrect inference. This issue is demonstrated with non-whitened particles, where the distributions of simulated and experimental particles do not overlap (Supplementary Figure 3).

We then applied a similar analysis on the experimental particles only, obtaining an insightful low-dimensional view of the entire experimental data set. Each point in Figure 4D corresponds to a single experimental image, colored according to the confidence with which the posterior maps it to a specific configuration. The confidence clearly describes a gradient correlating with the first reduced variable defining the plane (UMAP1). The confidence of the inference is high for particles on the left of the plot and gradually decreases going to the right. Particles on the left of the plot (low values of UMAP1) all contain a clearly visible copy of hemagglutinin in their center (Figure 4E). On the contrary, particles on the right (large values of UMAP1) do not contain any protein (Figure 4F). This observation shows that UMAP1 sorts particles according to how well hemagglutinin is visible. It quantifies how much structural information is contained in each particle.

A slender appendix in the 2D UMAP plot containing high-confidence points detaches from the distribution of points on the top of the plot. These particles clearly contain contaminants, e.g., unfolded proteins, structured ice, or other type of contamination (Figure 4G). Then, why does our inference lead to a high confidence? The posterior evaluated on these particles peaks at both extreme values of the prior range (Supplementary Figure 4). In other words, the posterior tells us that these particles are highly atypical and incompatible with all structures in our hypothesis ensemble, while they have low-resolution contrast that match some low-resolution template features. This observation is consistent with the appendix-like morphology of the region containing these particles. Typical particles are points with many neighbors. On the contrary, these particles are all on the border, making them very atypical. The analysis of the cryoSBI embedding is a powerful way of validating the inference and analyzing the experimental data set.

### 2.6 Template matching in a micrograph

The cryoSBI posterior can also match templates directly on a micrograph, bypassing the need to pick particles for the experimental data set. Figure 5A presents a hemagglutinin micrograph (3584*×* 3584 pixels) from the same dataset as discussed so far. For this micrograph, we employed a sliding window approach, using the trained cryoSBI posterior with a box size of 256*×*256 pixels. Evaluating many windows in parallel proved to be computationally efficient, achieving posterior evaluations for the entire micrograph in a couple of minutes. This allowed us to extract the posterior mean and width, which we then associated with the center of each box. Figure 5B illustrates the effectiveness of our method in identifying the cryo-EM reference structure from particles within the micrograph using the posterior mean. Examples of boxes with centers exhibiting a mean value close to the reference are shown in Figure 5C-E together with the posterior width. We note that because a convolutional neural network (which is translation equivariant) is used for the embedding, the exact particle position is not precisely determined. This results in a relatively wide range of pixels where the posterior mean closely matches the reference structure (red in Figure 5C-E). Analysis of the latent space combined with the shape of the cryoSBI posterior, could also facilitate the identification of outliers versus hemaggluting particles in the micrograph, as shown above for the picked particles.

**Figure 5:**
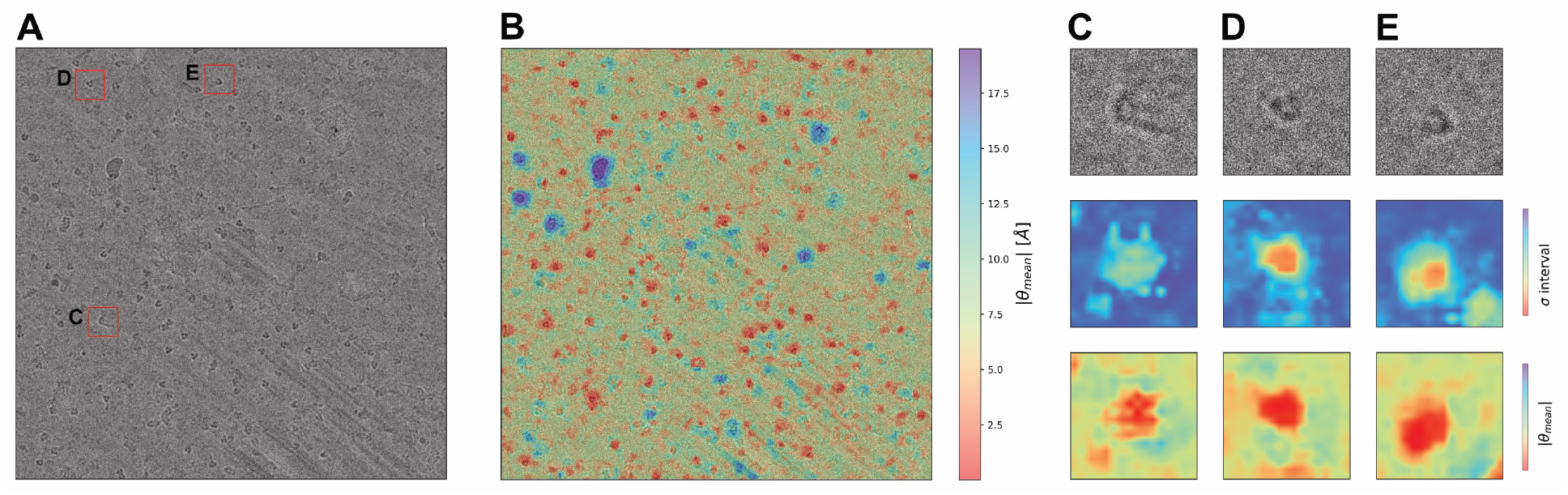
Evaluating the posterior learned by cryoSBI on a micrograph directly. A) Cropped micrograph (3824*×* 3824 pixels) from the hemagglutinin EMPIAR dataset 10026. B) Absolute Distance in *θ* between the posterior mean and the cryo-EM reference structure *θ* = 0 as a function of window position on the micrograph. C-E) Example windows (top) from the micrograph with a posterior mean close to the cryo-EM reference (bottom) and a small posterior width (middle).

## 3 Discussion

Reconstructing the conformational landscape of biomolecular systems from experiments is an outstanding challenge in molecular biology and biophysics. CryoSBI aims to overcome this problem by enabling Bayesian inference of biomolecular conformations from individual cryo-EM particle images. Given an initial structural ensemble hypothesis— a set of 3D templates—cryo-EM builds on simulation-based inference, a merger of physics-based simulations and probabilistic deep learning, to perform very fast amortized template matching in cryo-EM. Here, we show how this approach can produce accurate and precise inferences from noisy experimental cryo-EM particles.

The cryoSBI framework is based on Bayesian inference, which allows us to include prior knowledge and accurately assess uncertainties. Reconstructing the entire conformational landscape is a challenging inverse problem. To address this, we take a practical approach and divide the solution into two parts. Whether we use standard reconstruction or AI-based structure prediction, we can easily obtain one or a few reasonably accurate conformations. Then, molecular dynamics simulations, enhanced sampling, and artificial intelligence-methods can generate a hypothetical conformational ensemble from the initial few structures. This ensemble, which does not need to be entirely accurate but should provide a list of structural templates, becomes the prior for our inference.

The outcome of the inference is the posterior, a distribution describing the probability that a given particle contains a specific structural template. The cryoSBI posterior not only gives us information about the most probable conformation but also, crucially, provides an accurate statistical confidence interval given by the posterior width. The ability to go beyond point estimates is crucial to distinguish particles that provide useful structural information—characterized by a peaked posterior—from those for which the inference leads to very broad posterior distributions and are, therefore, not informative. In other words, some particles will be too noisy or originate from a specific pose such that a precise inference is not feasible, and the cryoSBI posterior will indicate this.

CryoSBI provides amortized inference, enabling the analysis of massive data sets. Given initial structural templates, cryoSBI trains an embedding and neural posterior density using simulated particles. Simulated template particles range over different conformations, poses, and other values of all the nuisance parameters associated with the image formation process. All these simulations are performed once upfront, after which we can train an embedding and a neural density estimator for the inference. The inference is a function only of the templates, marginalized over all other parameters, including the pose. The inference is amortized, i.e., we do not have to perform any optimization to solve the inference problem from scratch for each particle. We have only to evaluate the trained posterior, a forward pass evaluation of a neural network that takes of the order of milliseconds. Therefore, in contrast to most traditional and ML reconstruction frameworks, cryoSBI bypasses the pose and defocus estimate, resulting in an extremely fast inference. The amortized posterior enables the efficient application of cryoSBI to micrographs and data sets containing millions and billions of particles.

We still face significant challenges. In this work, we have demonstrated that cryoSBI can accurately infer large conformational changes in synthetic data and correctly identify conformations in experimental data. Our current focus is on expanding this framework to detect small conformational changes in large proteins and identify multiple alternative states. However, the most critical challenge is identifying and overcoming model misspecification. Any parametric inference is only as good as the model we assume to describe the underlying physical process. In our case, this is the structural ensemble that we use as a starting hypothesis and the simulator encoding the image formation process. Here, we have shown that the only real problem occurs when our ensemble is missing structures or when the simulator is missing features depicted in the experimental particles (e.g., Figure 4G). This instance leads to ‘hallucinations’, i.e., high-confidence inferences that are entirely wrong. However, we have also shown that the analysis of the latent space provides a powerful framework to diagnose model-misspecification. The accuracy of cryoSBI’s inference is only guaranteed if the distribution of points corresponding to the simulated particles largely contains the points corresponding to the experimental ones. This is equivalent to saying that the templates capture the underlying physical features in the cryo-EM data. A small overlap in the latent space would instead immediately reveal inadequate template simulations (e.g., Supplementary Figure 3). How to reduce model-misspecification by learning from data is an active field of research.

CryoSBI can improve current cryo-EM reconstruction pipelines. The posterior confidence can be used to classify and sort particles with sharp posteriors [46] and weight the particle contribution to the reconstruction. It could also be used to improve imaging conditions, such as the electron dose per frame, by monitoring the sharpness of posterior widths. An advantage of cryoSBI is that it provides a per-particle measurement plus an error, not suffering from orientational bias. Moreover, it can explain the relation between the conformational motion of interest and the projection direction (Figure 2B). Therefore, it could be combined with the ML heterogeneous reconstruction methods to sieve particles that do not provide information along the relevant conformational motion due to their projection direction. Even though we focused on inferring conformations in this work, cryoSBI can also be directly used to infer all other parameters involved in the image formation process, such as pose and defocus, setting priors, and ranges for cryo-EM reconstruction.

Cryo-EM has heavily relied on averaging particles. Even the state-of-the-art ML heterogeneous reconstruction methods rely on starting from a consensus volume and struggle with highly dynamic systems. CryoSBI provides a single-particle inference that can contribute to overcoming several problems, such as identifying rare conformations, structural intermediates (transition states), or studying highly flexible biomolecules by leveraging structural hypotheses from molecular simulations. The amortized inference could significantly speed up the recovery of free-energy landscapes or probability distributions from cryo-EM estimates [47, 23] by quickly comparing millions of particles to thousands of structures. Data-driven techniques applied to the latent space of experimental particles can lead to the discovery of new metastable states and the learning of the overall organization of the conformational landscape. Moreover, cryoSBI could identify particles *in situ* for visual proteomics within cellular contexts and study their environment-dependent properties by comparing the particles to the templates within the embedding space.

In summary, cryoSBI not only provides accurate structural inferences but also quantifies uncertainties through Bayesian posterior distributions. cryoSBI is a modular and flexible framework. The simulator, the embedding network, and the density estimator can readily integrate more sophisticated algorithms to overcome challenges emerging from complex datasets. Future work will focus on addressing model misspecification, enhance the simulator for more realistic scenarios, and expanding its capabilities to detect subtle conformational changes. These improvements will be vital for learning functional molecular conformational ensembles directly from experimental data in biologically relevant environments.

## 4 Methods and Materials

### 4.0.1 Cryo-EM image formation forward model

We used a standard forward model [41, 42] to simulate cryo-EM particles from a starting 3D molecular structure *X*. The model approximates the electron density *ρ*(*X*) as a Gaussian mixture model, comprising a number of Gaussian functions equivalent to the number of amino acids, all of equal amplitude. Each Gaussian is centered on the position of an amino-acid *C*_*α*_, with standard deviation sampled from a uniform distribution (see Biomolecular Systems section). The density *ρ*(*X*) is randomly rotated and translated. The rotation *R*_*q*_ is defined by a quaternion *q* and the translation by a vector ***τ***. The rotated density is then projected along the z-axis by the projection operator *P*_*z*_, using the weak-phase approximation, and then convolved with a point-spread function PSF, which models aberration and defocus of the microscope. In practice, we perform the convolution in Fourier space using the Fourier equivalent of the PSF, *i*.*e*., the Contrast Transfer Function defined as 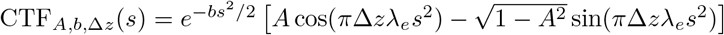 with the reciprocal radial component 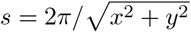, B-factor *b*, amplitude *A*, defocus Δ*z*, and electron wavelength *λ*_*e*_. Finally, we add Gaussian white noise to the images. The variance of the Gaussian noise is determined by the SNR and the amplitude of the signal, 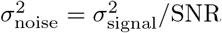, where *σ*_signal_ is the mean squared intensity of the image without noise within a circular mask of a given radius (for details see Biomolecular Systems section). The forward imaging model can be written as

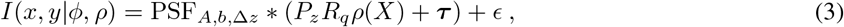

where the noise is drawn from a normal distribution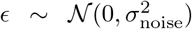, the imaging parameters are *ϕ* = *{q, τ, A, b*, Δ*z, σ*_noise_*}* and * denotes a convolution.

### 4.0.2 CryoSBI template simulations

In this paper, we parameterized a conformational ensemble with a 1D degree of freedom *θ*, defined as a function 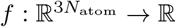, which maps an atomic structure *X* to a real number *θ* = *f* (*X*). We then selected a small number of molecular structures spanning *θ*, obtaining a discrete parameter *θ*_*i*_ containing our structural templates. The goal of cryoSBI is to learn the posterior probability *p*(*θ*_*i*_| *I*). To train cryoSBI, we run millions of forward model simulations of cryo-EM template images by using the forward model described above. We sample from the priors parameter distributions *p*(*θ*_*i*_, *ϕ*) = *p*(*θ*_*i*_)*p*(*ϕ*), where each *θ*_*i*_ selects a specific template, while *ϕ* contains the nuisance and imaging parameters, and generate a corresponding synthetic image.

An important aspect of cryoSBI is that it links (*θ*_*i*_, *ϕ*) to the image features, *S*(*I*), in a high-dimensional space (top middle Figure 1). The embedding network for the image featurization and a neural density estimator, a function that approximates the joint distribution of model parameters and features, are trained simultaneously. Details are provided below.

### 4.0.3 Priors

We chose a uniform prior *p*(*θ*_*i*_), giving each template equal *a priori* probability. While samples *θ*∼*p*(*θ*) from the prior are continuous, we used the closest representative template in *θ*_*i*_, with *min* |*θ*_*i*_− *θ*|. The priors for the imaging and nuisance parameters *p*(*ϕ*) were uniform within specific ranges (see details below for each system). Rotations were sampled from a uniform prior in SO(3) [48].

### 4.0.4 Embedding network

We used a modified ResNet-18 architecture [49] as embedding network *Sψ* (*I*) with parameters ψ to learn a compressed representation of the images *I*. We adapted the ResNet-18 to grayscale images and 256-dimensional output.

### 4.0.5 Learning the posterior

We used the Neural Posterior Estimation (NPE) algorithm to approximate the posterior distribution from synthetic particles [40]. NPE uses a neural network density estimator *q*_*φ*_ of parameters *φ* to approximate the posterior, *p*(*θ*| *I*) ≈ *q*_*φ*_(*θ*|*S*_*ψ*_ (*I*)). For each system, we created a large dataset of *N* synthetic particles *I*_*n*_∼ *p*(*I*|*θ*_*n*_, *ϕ*_*n*_) by drawing the prior conformations, *θ*_*n*_∼ *p*(*θ*), nuisance imaging parameters, *ϕ*_*n*_∼ *p*(*ϕ*), and then running a total of *N* forward model cryo-EM template simulations, with *n* = 1, …, *N*. We then trained jointly the embedding network and the density estimator by maximizing the average log-likelihood of the posterior probability under the training samples

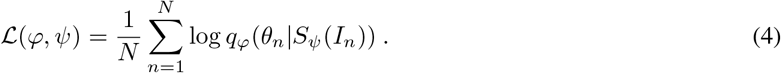

We used a Neural Spline Flow (NSF) [50] as density estimator, containing 5 transformation stages. The neural network in each transformation stage contained 12 layers. The first ten layers had 256 hidden nodes, while the last two had 128 and 64 hidden nodes, respectively. We trained the network using an AdamW optimizer [51], with a learning rate of 0.0003, gradient clipping with a maximum norm of 5, and a batch size of 256. Because the forward model simulation is inexpensive, we generated synthetic images during training on demand and did not store the training set. Therefore, each batch of images was newly generated, allowing training until convergence and preventing overfitting.

### 4.0.6 Amortized posterior evaluation

To evaluate the trained posterior *p*(*θ*| *I*^*′*^)≈ *q*_*φ*_(*θ*|*S*_*ψ*_(*I*^*′*^)) conditioned on an observed image *I*^*′*^, we generated samples by drawing directly from the neural posterior using the trained density estimator and embedding network [49]. The process includes drawing samples from the base Gaussian distribution of the NSF, transforming them using the normalizing flow, and simultaneously computing the latent representation for the image of interest that will condition the posterior. This entire procedure is automatically parallelized through batch computations in PyTorch. Typically, we generated around 20,000 posterior samples per image to estimate the continuous posterior density.

### 4.0.7 Latent-space analysis

To identify model misspecification, we compared the latent representations of the embedding network of experimental and simulated particles. We evaluated the similarity between these sets of points in the latent space in two ways. The first is a qualitative visual inspection relying on the dimensionality reduction technique UMAP [52]. We concatenated the latent representations (256 dimensions) from the simulated and experimental particles, and performed a UMAP analysis to identify a 2D projection. We inspected the 2D UMAP to make sure that simulated and experimental particles overlapped.

In a second, more quantitative way, we compared the distributions of the latent representation of experimental and simulated particles using the maximum mean discrepancy (MMD) metric. MMD is a standard metric to statistically test whether two independent sets of samples come from two different distributions [53], and is often used in the SBI community to test model misspecification [54].

Let *S*_1_ = *{s*_1_*}* and *S*_2_ = *{s*_2_*}* be two datasets containing the 256-dimensional latent representations. For the MMD, we used a Gaussian kernel with Euclidean distances in the embedding space, so that for a pair of *s*_1_, *s*_2_, the kernel is *k*_*ϵ*_(*s*_1_, *s*_2_) = exp(−||*s*_1_− *s*_2_|| ^2^*/ϵ*) where||. || ^2^ is the *l*2-norm. We choose the bandwidth *ϵ* as the median of pairwise squared distances between the datasets *S*_1_ and *S*_2_.

### 4.1 Biomolecular systems

#### 4.1.1 Hsp90

We used Hsp90 as a first system to validate cryoSBI by having ground truth synthetic data [41, 42]. Hsp90’s conformational landscape can be described by the relative motions of its chains, A and B. Here, we focus on the opening of chain B depicted by *θ* in Figure 2A, and measured it as the RMSD with respect to the closed structure. We built a set of 20 templates *θ*_*i*_, with *θ*_*i*+1_ −*θ*_*i*_ = 2^°^ angle difference in chain B’s displacement. We produced synthetic particles with image size 128*×*128 pixels, and a pixel size of 1.5 Å. Each residue of Hsp90 was represented by a Gaussian density with a standard deviation whose prior ranged between 0.5 and 5 Å. The SNR prior ranged from 0.5 to 0.001. The signal of the images without noise was determined by a circular mask with a radius of 64 pixels. For the CTF, the defocus prior was sampled uniformly between 0.5 and 2 *µ*m, the amplitude A was constant at 0.1, and the B-factor was sampled uniformly between 1 and 100 Å^2^. To learn an amortized posterior, we trained the NSF density estimator for 150 epochs with 2000 batches per epoch. To compare cryoSBI to an explicit-likelihood method, we used the BioEM software [31] with the same parameter ranges, and sampled orientations with a uniform grid of 36864 quaternions on SO(3) [55].

#### 4.1.2 Apoferritin

We used apoferritin experimental images [44] available at EMPIAR 10026. We started from the reconstructed high-resolution cryo-EM structure (PDB id: 4v1w) and generated a set of templates by building normal modes using ProDy [56]. To avoid multimodal posteriors due to symmetry, we concatenated two normal mode displacements, modes 1 and 10, with maximum displacements of 12.0 Å (left and right side of Figure 3A), from the reconstructed cryo-EM structure. We defined *θ* = *γ* RMSD where *γ* =− 1 if the structure *x* belongs to mode 1 and *γ* = 1 if the structure belongs to mode 2, calculating the RMSD is with respect to the cryo-EM structure. We obtained 51 templates *θ*_*i*_ with a *θ*_*i*+1_− *θ*_*i*_ = 0.48Å. We simulated synthetic cryo-EM particles with an image size of 132*×*132 pixels and a pixel size of 1.346 Å, equivalent to the experimental particles. Each residue of apoferritin was represented by a Gaussian density centered at the *C*_*α*_ position, with standard deviation uniformly sampled between 0.5 and 5 Å. The SNR ranged from 0.1 to 0.001. The signal of the simulated images without noise was determined by using a circular mask with a radius of 66 pixels. For the CTF, the defocus was sampled uniformly between 1.0 and 4.0 *µ*m, the amplitude was constant at 0.1, and the B-factor was sampled uniformly between 1 and 100 Å^2^. The orientations were sampled uniformly in SO(3). To learn an amortized posterior, we trained the NSF density estimator for 100 epochs with 1600 batches per epoch. The experimental particles were whitened using the Python implementation of ASPIRE [57].

#### 4.1.3 Hemagglutinin

We used the hemagglutinin experimental images available at EMPIAR 10532 [45]. A subset of these particles were used to reconstruct a structure at around 3Å resolution (PDB id: 6wxb). Similarly to apoferritin, we used cryoSBI and normal mode displacements to test if the cryo-EM structure can be inferred from each individual particle. The normal mode trajectory was generated using ProDy [56]. To avoid multimodal posteriors due to symmetry, we concatenated two normal mode displacements mode 1 and 2 (left and right side of Figure 4A-bottom) from the cryo-EM structure. The maximum displacement for mode one is 20 Å and for mode two 8 Å. We defined *θ* = *γ* RMSD where *γ* =−1 and 1 for structures from mode 1 and 2, respectively. Normal mode structures were mapped along *θ* with a 0.4 Å interval. To train cryoSBI, we simulated cryo-EM templates with an image size of 128*×*128 pixels and a pixel size of 2.06 Å. Each residue of hemagglutinin was represented by a Gaussian density centered at the *C*_*α*_ position with standard deviation uniformly sampled between 0.5 and 5 Å. The SNR ranged from 0.1 to 0.001. The signal of the simulated images without noise was determined by a using circular mask with a radius of 64 pixels. For the CTF, the defocus was uniformly sampled between 0.5 and 5 *µ*m, the amplitude was constant set to 0.1, and the B-factor was set to 1 Å ^2^. The orientations were sampled uniformly from SO(3). We applied a low pass filter cutting frequencies higher than 10Å. The low pass filter was applied in order to reduce artifacts from the background. We trained the neural network density estimator for 450 epochs, each containing 800 batches. To test the experimental particles with cryoSBI, we first downsampled them to match the pixel and box sizes of the simulated images. Then, we whitened them using the Python implementation of ASPIRE [57]. Whitening is necessary to avoid a noise-model misspecification, which can be assessed by comparing the projection of the simulated templates and the experimental in the latent space of the embedding (Supplementary Figure 3B).

##### Latent-space analysis of hemagglutinin

We used the MMD to compare the entire set of simulated templates with SNR ranging between 0.001 and 0.1 to subsets of simulated templates with different SNR ranges, and to sets of raw and whitened experimental particles (Supplementary Figure 3A). Due to the large dataset sizes of 271558 particles, we approximated the MMD by bootstrapping with random subsets of size 10^4^ from each set, and then took the average MMD over 100 trials.

##### Hemagglutinin micrograph analysis

We analyzed a cropped micrograph (3824*×*3824 pixels) using a 256×256 pixel large sliding window, moving it in steps of 16 pixels. This resulted in a total of 58081 (241*×*241) windows. We whitened each window using the ASPIRE Python implementation [57] and downsampled it to 128×128 pixels. In each window, we evaluated the same posterior learned for the hemagglutinin single particles. To visualize different posterior properties as a function of the crop center position, we upscaled the 241*×*241 matrix to the size of the micrograph.

##### Molecular dynamics simulation of hemagglutinin

We ran a molecular dynamics simulation of hemagglutinin using the Charmm36m [58] forcefield to estimate the fluctuations around its folded conformation. We started from the cryo-EM structure PDB: 6WXB. We added 24 missing residues and solvated the structure with TIP3P water with 0.15M of NaCl. We minimised the energy of the systems with 5000 steps of the steepest-descent algorithm. Afterwards, we performed a 125 ps long equilibration in the NVT ensemble using the Nose-Hoover thermostat set at a reference temperature of 310 K. We simulated the systems in the NPT ensemble for 557.75 ns, keeping a reference temperature of 310 K (Nose-Hoover thermostat with *τ* t = 1 ps) and a reference pressure of 1 bar (Parrinello-Rahman barostat with *τ* p = 5 ps). We used a radius-cut-off = 1.2 nm for the Van der Waals (Verlet) and the Coulomb forces (Particle Mesh Ewald). We used a simulation timestep of 2 fs. We computed the pairwise RMSD for residues which have a RMSF of 2.0 Å or lower within the trajectory to estimate the expected fluctuations. We calculated a histogram from the pairwise RMSD values and selected the value at the 68% quantile, corresponding to approx. 2.0 Å (Supplementary Figure 1). Therefore, we expect that in 68% of cases, the difference between structures will be not more than 2Å (Figure 4B dotted lines around dashed line).

### 4.2 Code Availability

The code is available at GitHub https://github.com/flatironinstitute/cryoSBI and is based on LAMPE [59], a PyTorch implementation for simulation-based inference. Data and scripts necessary to reproduce all the results in this paper are freely accessible at the Zenodo repository https://zenodo.org/records/12608562.

## 5 Acknowledgments

The Flatiron Institute is a division of the Simons Foundation. E.D.I acknowledges IRCCS Humanitas Research Hospital for financial support. L.D. and R.C. acknowledge the support of Goethe University Frankfurt, the Frankfurt Institute of Advanced Studies, the LOEWE Center for Multiscale Modelling in Life Sciences of the state of Hesse, the CRC 1507: Membrane-associated Protein Assemblies, Machineries, and Supercomplexes (P09), and the International Max Planck Research School on Cellular Biophysics, as well as computational resources and support from the Center for Scientific Computing of the Goethe University and the Jülich Supercomputing Centre. D.S.S, L.D., and R.C. thank the Flatiron Institute for hospitality while a portion this research was carried out.

## A Estimating structural flexibility of hemagglutinin with molecular dynamics simulations

**Supplementary Figure 1:**
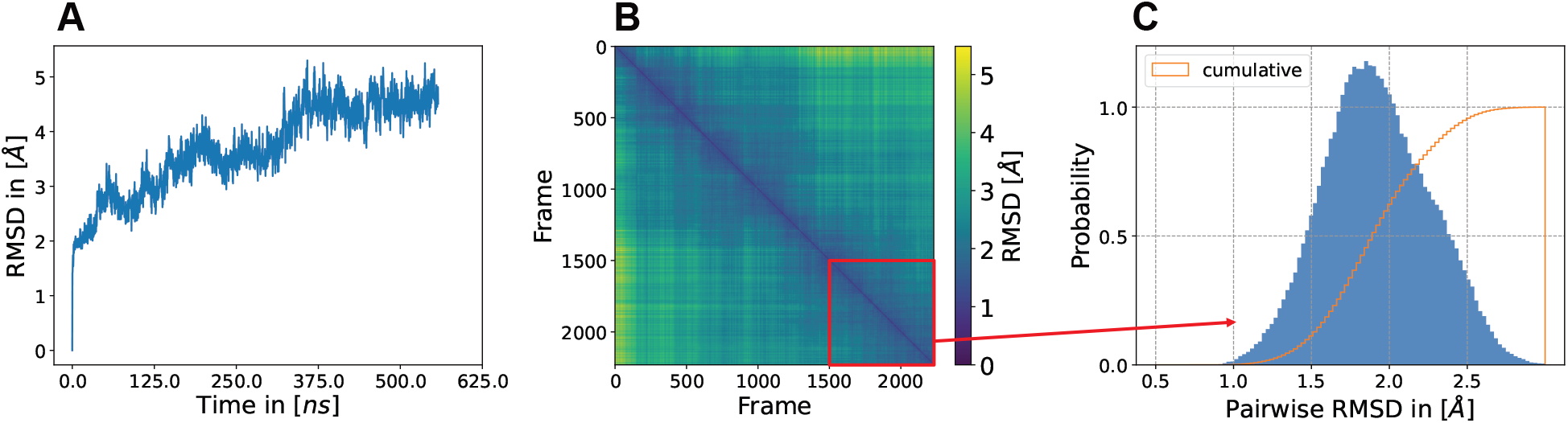
MD simulations of hemagglutinin starting from the cryo-EM reference structure. A) RMSD calculated along the trajectory w.r.t. the cryo-EM reference structure. B) Pairwise RMSD of all trajectory trajectory frames. C) Histogram of the pairwise RMSD values for the last 182 ns of the trajectory. The total simulation time was 557 ns.

## B UMAP analysis for hemagglutinin

**Supplementary Figure 2:**
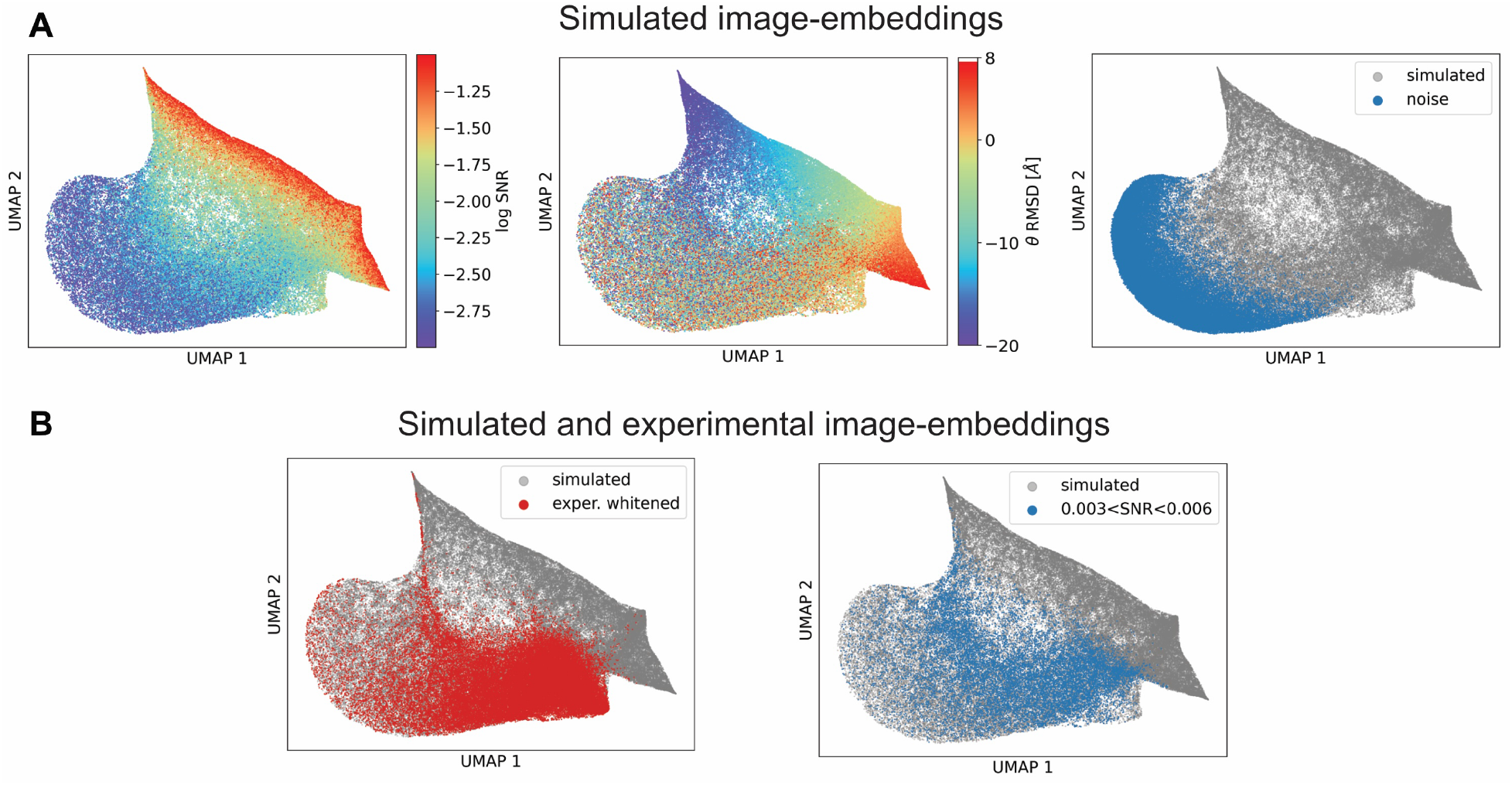
UMAP comparison of simulated templates and experimental particles’ embedding of hemagglutinin. UMAP was computed from the latent space of 10^5^ simulated templates, 50, 000 simulated templates with no particle present (“noise”), and a random subset of 50, 000 whitened experimental particles. A) The UMAP of simulated templates is colored by log-SNR (left plot) and by *θ* (middle plot). The UMAP of the pure noise images (blue) is also shown with the simulated images (gray) (right plot). B) Experimental whitened particles have a similar MMD to the set of simulated templates with SNR range ∈ [0.003,0.006] (Supplementary Figure 3A). We compare these sets using UMAP. The UMAP of whitened experimental particles depicted in red (left plot) and simulated templates with SNR range ∈ [0.003,0.006] in blue (right plot) with simulated templates in the full SNR range in gray.

## C Maximum Mean Discrepancy model misspecification for hemagglutinin

**Supplementary Figure 3:**
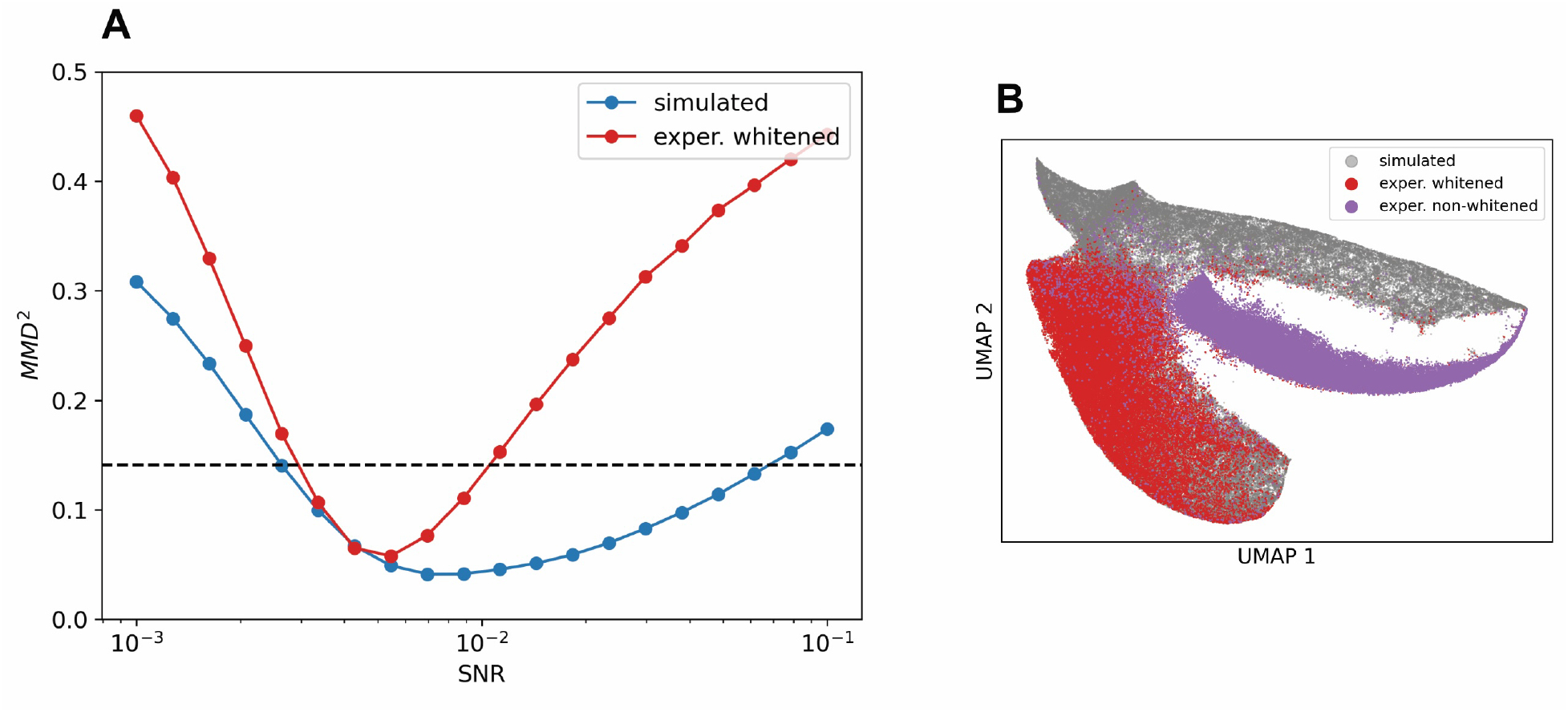
A) Maximum Mean Discrepancy (MMD) analysis for hemagglutinin. We use MMD to compare the embeddings of different subsets of templates and experimental particles in the 256-dimensional latent space. We show the MMD of 10^5^ simulated templates at a *fixed* SNR_*i*_ (x-axis) with respect to 10^5^ simulated templates with SNR in the full range ∈ [0.001, 0.1] (blue), to the dataset of whitened experimental images (red), and to the dataset of non-whitened experimental images (purple). The black dotted line is the MMD between simulated templates in the full SNR range ∈ [0.001, 0.1] and the whitened experimental dataset. We highlight the SNR region where the MMD whitened experimental particles closely matches the simulated templates. B) 2D UMAP computed from the latent space of 10^5^ simulated templates (gray), a random subset of 50, 000 whitened experimental particles (red), and the same particles but non-whitened (purple).

## D Posteriors of misspecified particles

**Supplementary Figure 4:**
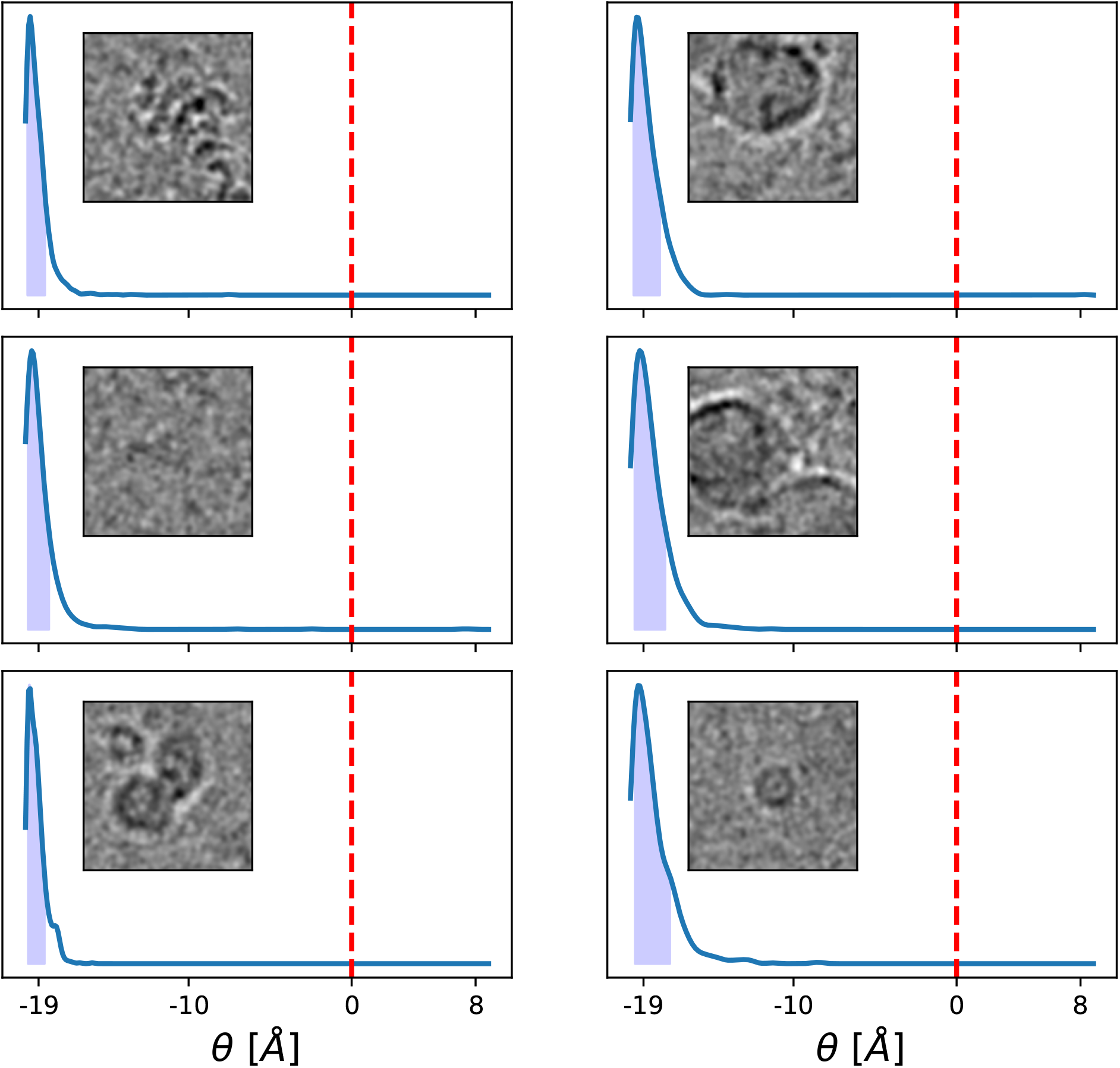
CryoSBI posteriors for misspecified hemagglutinin particle images. The particles are low-pass filtered for better visibility. The conformational change along *θ* is described using normal mode analysis, where *θ* = *γ* RMSD from the reference cryo-EM structure (red dashed line), with *γ* equal to -1 or 1 for normal mode 1 and 2, respectively.

